# Exploring the Effects of Ecological Parameters on the Spatial Structure of Genealogies

**DOI:** 10.1101/2023.03.27.534388

**Authors:** Mariadaria K. Ianni-Ravn, Martin Petr, Fernando Racimo

## Abstract

Geographic space is a fundamental dimension of evolutionary change, determining how individuals disperse and interact with each other. Consequently, space has an important influence on the structure of genealogies and the distribution of genetic variants over time. Recently, the development of highly flexible simulation tools and computational methods for genealogical inference has greatly increased the potential for incorporating space into models of population genetic variation. It is now possible to explore how spatial ecological parameters can influence the distribution of genetic variation among individuals in unprecedented detail. In this study, we explore the effects of three specific parameters (the dispersal distance, competition distance and mate choice distance) on the spatial structure of genealogies. We carry out a series of *in silico* experiments using forwards-in-time simulations to determine how these parameters influence the distance between closely- and distantly-related individuals. We also assess the accuracy of the maximum likelihood estimation of the dispersal distance in a Gaussian model of dispersal from tree-sequence data, and highlight how it is affected by realistic factors such as finite habitat size and limited data. We find overall that the scale of mate choice in particular has marked patterns on short and long terms patterns of dispersal, as well as on the positions of individuals within a habitat. Our results showcase the potential for linking phylogeography, population genetics and ecology, in order to answer fundamental questions about the nature of spatial interactions across a landscape.

## 2 Introduction

From nutrient-fixing bacteria in the digestive system, to pollen carried on the legs of bees, all living organisms must deal with the particularities of the range that they inhabit. At each generation, individuals tend to disperse from their parents, often carrying their genes across great geographic distances. Geographic space is also a major determinant of mate choice and competition for finite resources such as food and water, which can, in turn, influence how genetic relatedness decays as a function of the distance between individuals (Wright 1943). The connection between genetic differentiation and geography has indeed been the focus of numerous theoretical models (for instance François Rousset 1997; Hardy and Vekemans 1999; B. Charlesworth, D. Charlesworth, and Barton 2003; Robledo-Arnuncio and Francois Rousset 2010) and empirical studies (Sexton, Hangartner, and Hoffmann 2014; Jenkins et al. 2010; Aguillon et al. 2017). Overall, it is now clear that genetic data can hold information about the geographic distribution of individuals in the past (Novembre et al. 2008; Aguillon et al. 2017).

Biologists often seek to understand the rate at which individuals of a given species move across space. One way to approach this problem is by focusing on the “dispersal distribution”: a probability distribution over the parent-offspring distance (Kot, Lewis, and Driessche 1996) i.e. how far away a particular offspring mates compared to its birthplace. The shape of the dispersal distribution for different species has been of great interest in ecology. In particular, long-distance dispersal is predicted to strongly affect patterns of relatedness across a species (T. B. Smith and Weissman 2020), as well as population genetic processes such as allele surfing (Paulose and Hallatschek 2020) and ecological phenomena including the spread of invasive species and host-parasitoid interactions (McCann et al. 2000; Clark 1998).

The dispersal distribution is often summarized via a “dispersal distance” parameter, *σ*, which predicts how far away an offspring tends to be from its parents. More precisely, *σ* should be seen as an “effective” dispersal parameter, which absorbs several stages of mate choice and parent or offspring migration, to predict how far a successfully reproducing offspring moves from its birth location (Bradburd and Ralph 2019; C. C. Smith et al. 2023). Over multiple generations — for example, over branches in a phylogeny — this determines the speed at which two lineages move away from one another after descending from a common ancestor (Francois Rousset 2001). It is known that the rate of geographic dispersal affects genetic variation (B. Charlesworth, D. Charlesworth, and Barton 2003). Conversely, it is possible to learn *σ* from genotype data with some accuracy (François Rousset 1997; Ringbauer, Coop, and Barton 2017; C. C. Smith et al. 2023).

One way to estimate the parameters of the dispersal distribution in a real population is to track the exact locations of all individuals in a pedigree. However, this is often difficult or expensive (Cayuela et al. 2018). While non-recombining genetic sequences can be easily recorded in a genealogy or coalescent tree (Miles et al. 2009; Markov et al. 2009; Castillo et al. 2011), the full history of recombining genomes cannot. Instead, this may be represented as a network, known as the Ancestral Recombination Graph (ARG) (Hudson et al. 1990; Griffiths and Marjoram 1996; Griffiths and Marjoram 1997), which fully encodes the history of coalescence and recombination of a set of sampled genomes. An alternative representation is an ordered set of coalescent trees, each describing the history of a section of the genome in the samples (a “tree sequence”, Kelleher, Wong, et al. 2019). Adjacent genealogies are separated by recombination events, and tend to be more highly correlated than those representing distant genomic tracts. A tree sequence can encode the full ARG, if it contains certain annotations (Rasmussen et al. 2014).

Genome-wide tree sequences are an ideal object on which to perform phylogeographic inference, and are already beginning to be used for such analyses (for example, Wohns et al. 2022). Recent computational developments have made it tractable to approximately infer tree sequences for a given genome panel (Kelleher, Wong, et al. 2019; Speidel et al. 2019; Hubisz and Siepel 2020). However, both estimating and working with full tree sequences comes with substantial computational burden. One approach to this problem, which has been used in recent work (Osmond and Coop 2021), is to “thin” the full sequence of trees covering the entire chromosome into a set of approximately independent genealogies. Although these genealogies do not wholly capture the complexity of the full tree sequence, we believe that the insights obtained from them are an important basis for understanding how spatial dispersal affects recombining genomes.

In recent years, new software for generating spatially explicit, forwards-in-time simulations have enabled researchers to explore genetic variation under a wide range of population histories. The recently developed software *slendr* (Petr et al. 2022), which uses the powerful software *SLiM* as one of its simulation engines (Haller and Messer 2019; Haller, Galloway, et al. 2019) provides a particularly approachable way to model, visualize and simulate mate choice, dispersal and spatial interactions in continuous space. These simulators can bridge the gap between a lack of theoretical results and the desire to build realistic spatial models of species.

Two types of interactions which people often use to model populations in geographic space are mate choice and competition for resources. Both of these can be understood via a distance parameter. The mate choice distance controls the scale at which individuals tend to find each other to produce an offspring. The competition distance determines how far individuals can be separated for them to compete for resources. The effects of these parameters on dispersal and genetic diversity have not been the main focus on previous studies. However, there is some evidence from simulations that the scale of mating has more impact on effective dispersal than that of competition (C. C. Smith et al. 2023). The lack of work in this area is particularly troublesome for any users of forwards-in-time simulators such as SLiM, where they are required to specify these dynamics explicitly.

Motivated by these issues, here we set out to understand properties of geographically annotated sequences of genealogies along a genome, using a simulation-based approach. We leverage *slendr* to carry out forwards-in-time simulations with non-overlapping generations, and study how ecological parameters affect the spatial distribution of individuals, and the structure of genealogies relating them over time.

First, we explore the effects of varying the mode and scale of mating and dispersal on the realised distances between parents and their offspring. We show that, in some cases, these distances closely match their theoretical distribution. We find that the scale of mate choice is an important determinant of the shape of dispersal distributions and the overall rate of dispersal. Then, we illustrate a case in which the realised distribution closely matches a theoretical model which explicitly includes the radius of mate choice. Finally, we test the efficacy of a maximum likelihood estimator of the mean distance between parent and offspring, using distances recorded in the branches of a phylogeny under a commonly used Gaussian mode of dispersal.

Our work serves to show that a sound understanding of the geographic parameters of a species, with respect to the dispersal distribution and to ecological factors (such as competition for resources and mate choice), is key to carrying out reliable phylogeographic inference in real populations.

## 3 Results

### 3.1 Dispersal patterns in spatially-tagged genealogies

We were interested in learning the relationship between the observed parent-offspring distances in a (perfectly inferred) genealogy and the underlying dispersal function in a population. In our simulations, a dispersal function (*DF*) and its scale parameter, *σ*, determine how the simulator decides where to place offspring compared to the gestating parent (p1). More details on this process are described in the Methods section 5.1.1, and a schematic of these mechanics is shown in Fig. 1. The range of *DF* s and their parametrization are summarised in Table 1 and plotted in Fig. 2. Although the interpretation of *σ* with respect to the *DF* varied for each distribution, our parametrization was such that increasing *σ* increased the variance of parent-offspring displacement. In essence, the larger *σ*, the further an offspring tends to be from its parents and the faster genetic lineages spread across the habitat.

**Table 1:**
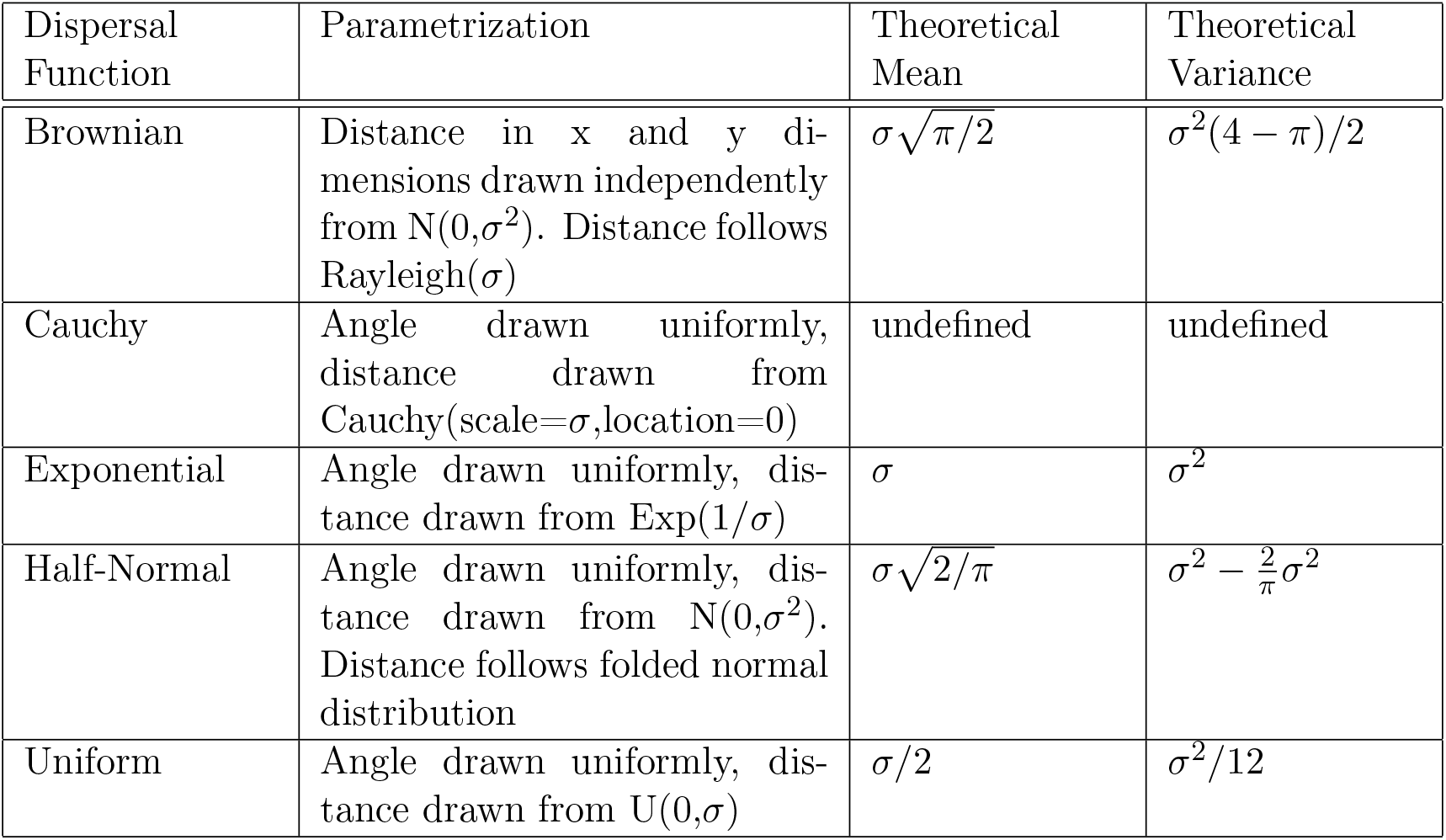
The parametrization of parent-offspring distances via the dispersal distance. We parametrized the dispersal distribution through a parameter *σ*, such that the theoretical variance increased with *σ*^2^, and the mean with *σ* (this does not apply to the Cauchy distribution, which has undefined mean and variance; here, *σ* was the scale parameter). Further details are given in Methods section 5.1.1.

| Dispersal Function | Parametrization | Theoretical Mean | Theoretical Variance |
| --- | --- | --- | --- |
| Brownian | Distance in x and y dimensions drawn independently from $N(0, \sigma^2)$. Distance follows $\text{Rayleigh}(\sigma)$ | $\sigma\sqrt{\pi/2}$ | $\sigma^2(4 - \pi)/2$ |
| Cauchy | Angle drawn uniformly, distance drawn from $\text{Cauchy}(\text{scale}=\sigma, \text{location}=0)$ | undefined | undefined |
| Exponential | Angle drawn uniformly, distance drawn from $\text{Exp}(1/\sigma)$ | $\sigma$ | $\sigma^2$ |
| Half-Normal | Angle drawn uniformly, distance drawn from $N(0, \sigma^2)$. Distance follows folded normal distribution | $\sigma\sqrt{2/\pi}$ | $\sigma^2 - \frac{2}{\pi}\sigma^2$ |
| Uniform | Angle drawn uniformly, distance drawn from $U(0, \sigma)$ | $\sigma/2$ | $\sigma^2/12$ |

**Figure 1:**
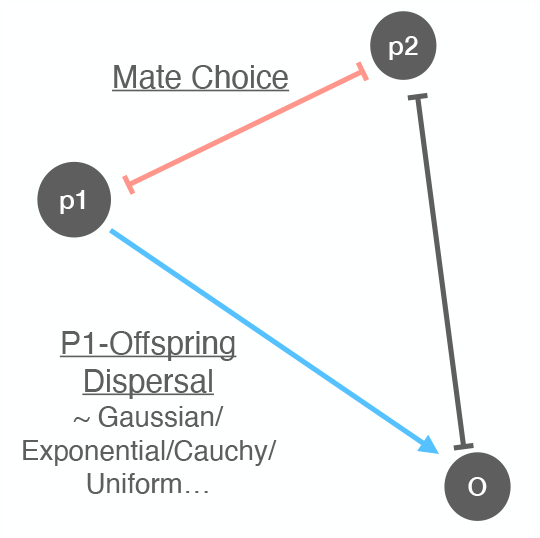
The mechanics of dispersal in our simulations. In our forwards-in-time simulations, two parents *p*1 and *p*2 are chosen. The distance between them (red line) must be less than the user-specified mating distance. The offspring (*o*) is then dispersed from *p*1 (blue line) according to a specified mode of dispersal parametrized via a dispersal function (*DF*) and distance (which we call *σ*). These mechanics imply that a given one-generation dispersal may either be a direct observation of a draw from the *DF* (the *p*1 − *o* distance, blue line) or it may be a composite of mate choice and dispersal (the *p*2 − *o* distance, grey line).

**Figure 2:**
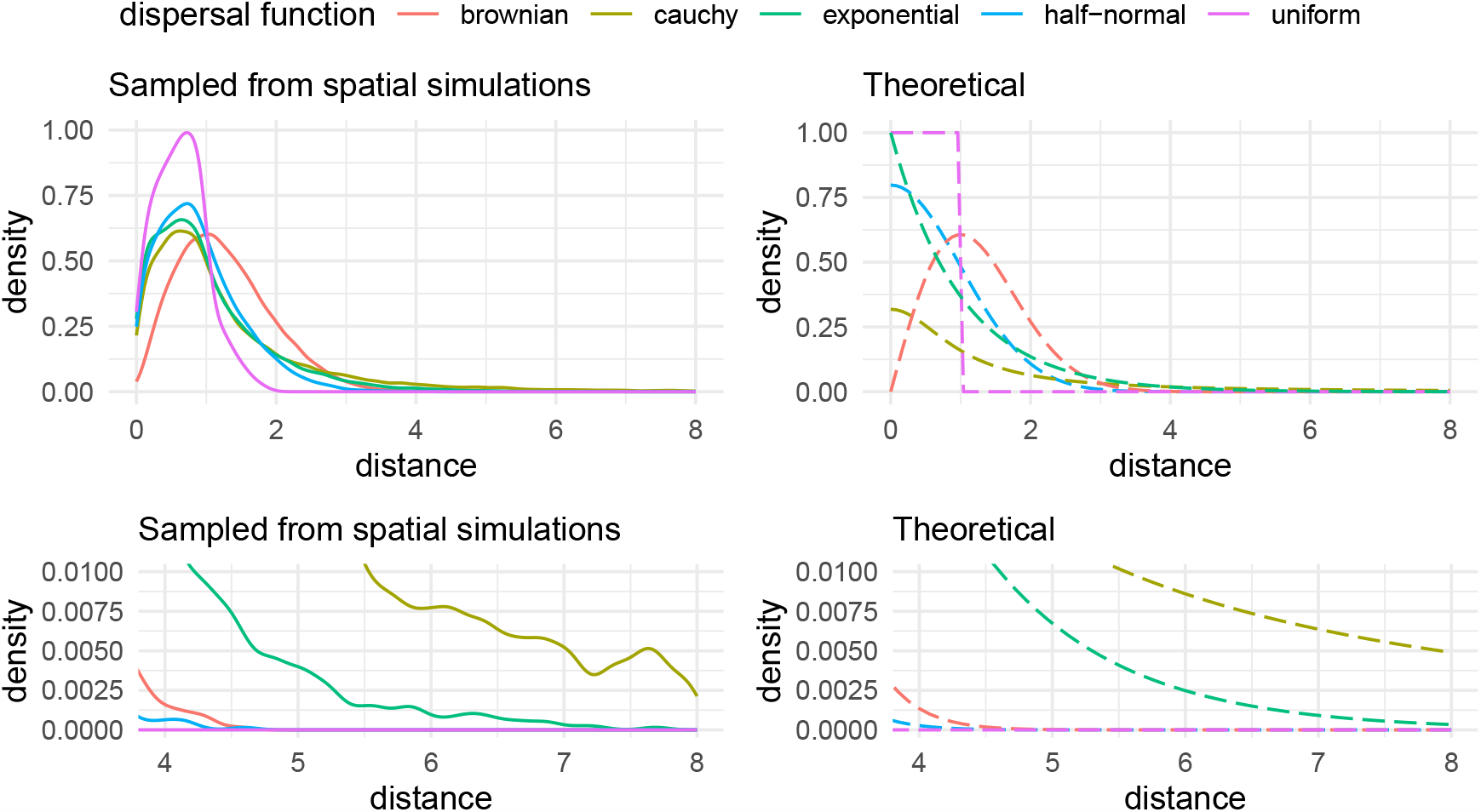
Distributions of parent-offspring distances reflect the underlying dispersal function. The left panel shows the empirical distribution of effective parent-offspring distances drawn from the forwards-in-time spatial simulations, while the right one shows the PDF of the corresponding dispersal distributions. The effective distances are affected by the dispersal distribution, as well as competition and mate choice. Bottom: a zoom-in on the tails of curves; the height of the tails of the distributions corresponded to those of the corresponding dispersal functions, with the Cauchy having the most heavy tail, followed by the exponential, Brownian, half-normal and then uniform. The mating distance competition distances were both 1 unit.

Two other important parameters in our simulations are the competition and mating distances. The competition distance serves to parametrize competition for resources within a neighbourhood. Essentially, the simulator counts the number of neighbours an individual has within a radius of the competition distance and down-scales their fitness proportionally to this number (see Methods section 5.1.1). The mate choice radius, or mating distance, determines the maximum radius within which a parent can choose a mate. In *slendr* V0.5.1, mates are chosen uniformly at random from within this distance.

We simulated a single, non-recombining locus in a population of 100 individuals in a habitat of size 50 *×* 50 units. We used a range of dispersal functions and *σ* values, and also varied the mating and competition distances. After 50 generations, we sampled all individuals and reconstructed the genealogy connecting them. In these genealogies, we stored all individuals, rather than coalescent nodes only (this corresponds to a tskit “unsimplified” tree), so that we could observe dispersal at every generation. For each condition, we ran 20 replicates. We will call the distribution of realised parent-offspring distances in these trees the 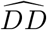 (empirical distance distribution).

We compared parent-offspring distances sampled from the simulations (the 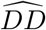) to the theoretical probability distributions from which p1-offspring distances were drawn (the *DF*). The shape of the 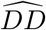 tended to mirror that of the *DF* (Fig. 2). For example, when parameterizing the *DF* as Cauchy, we observed a higher frequency of long 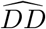 dispersal values, compared to other *DF* distributions, when the parameter *σ* was kept constant. This is consistent with the heavy tail of the theoretical Cauchy distribution, compared to other distributions (uniform, half-normal, exponential or Rayleigh).

There was not a perfect correspondence between 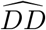 and *DF*, as the other ecological parameters (namely competition distance and mate choice radius) in the simulation also influenced the realized distance between parent and offspring. We quantified this effect of these parameters on effective dispersal by measuring the excess variance of the 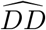, compared to the *DF* (Fig. 4). Increasing the mating distance caused the 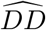 to accumulate much excess variance, and the 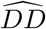 to acquire a flat “shoulder”, which we model in the next section (section 3.2).

**Figure 3:**
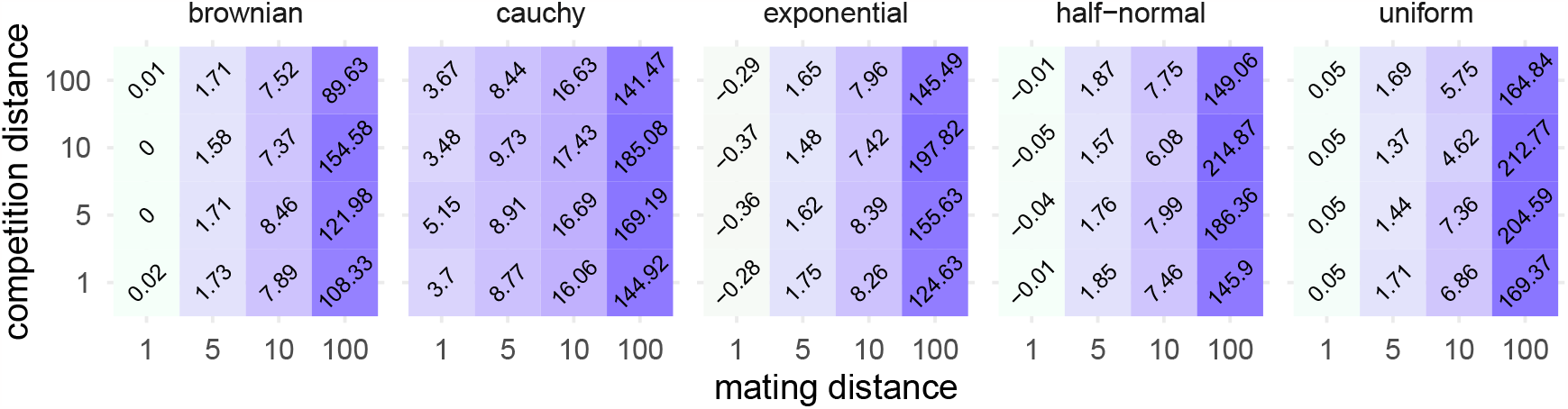
Quantifying the effect of mate choice and competition radius on realized parent-offspring distances. Each tile shows the excess variance of the empirical dispersal distribution compared to the theoretical one — as given by Table 1. Since the Cauchy distribution has undefined variance, the excess is relative to zero. Increasing the competition distance tended to have relatively little effect on the variance of parent-offspring distances, but altering the scale of mate choice had a very strong effect.

**Figure 4:**
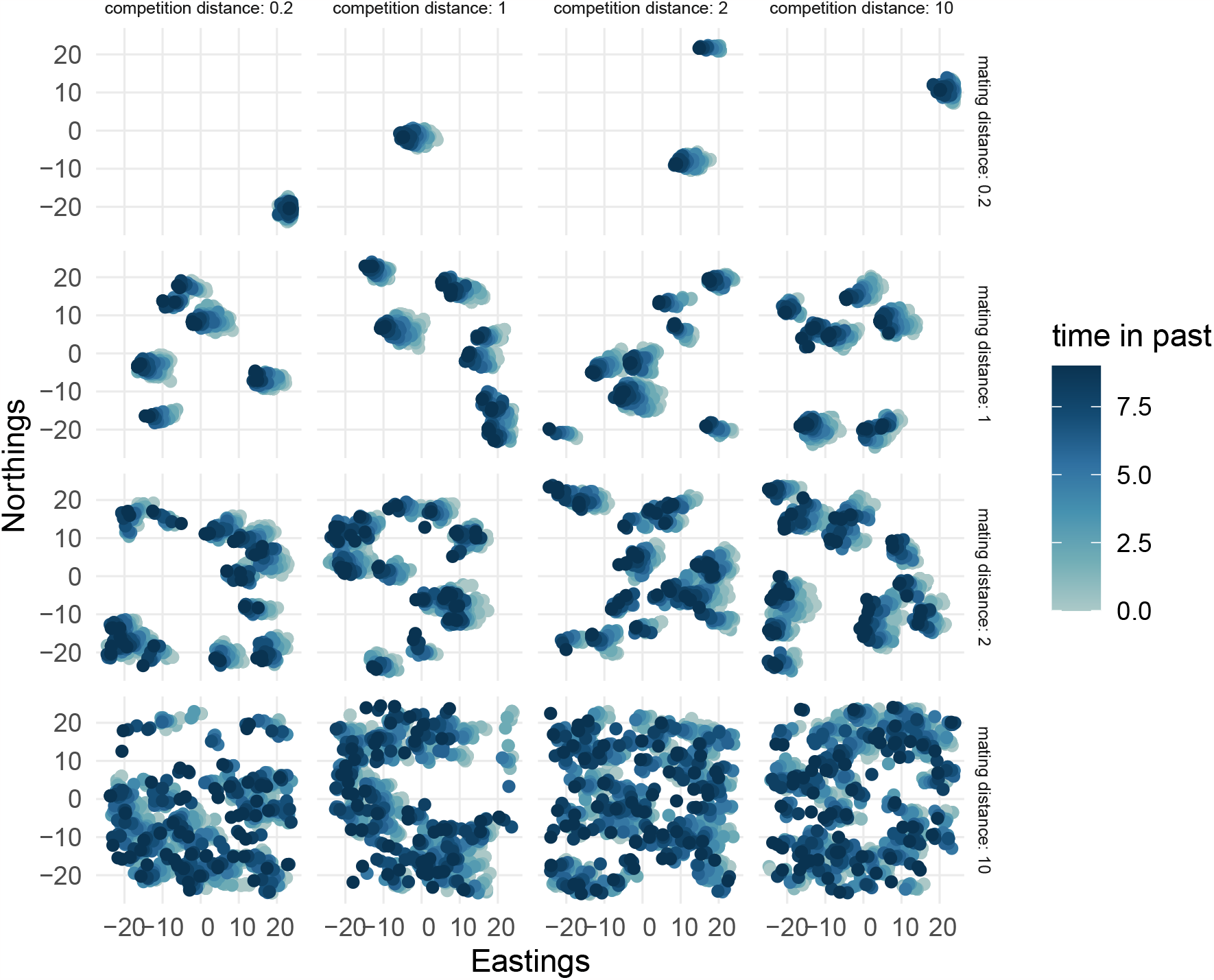
The effects of the mating and competition distance on the placement of individuals in a population. The positions of individuals present in the genealogy connecting 200 sampled individuals (in the unsimplified tree), over 10 generations are shown and coloured by time. When the mating distance was small (top row), we observed a strong clumping behaviour. As we increased the competition and mating distances, the clustering behaviour was alleviated. In these simulations, the mode of dispersal was Brownian and *σ* was 1 unit.

In contrast, varying the competition distance had a weaker effect on excess variance (the difference between the variance of the 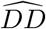 and the *DF*, brought about by mate choice and competition). Excess variance tended to increase with competition distance; however, when the competition distance was 100 (twice the width of the habitat), the effect on the excess variance was small. This was expected, since a radius of 100 spans the entire 50 *×* 50 habitat we simulated, and therefore is equivalent to no competition at all (since every individual’s fitness is down-scaled by the same factor, the total population size, leading to equal relative fitness across the population — for details, see Methods section 5.1).

Overall, the relationship between the theoretical and realized parent-offspring distributions — under varying competition and, in particular, mating distances — suggests that these ecological parameters may be important determinants of the scale of dispersal of individuals in the wild.

To further investigate the nature of the effects of mating and competition, we examined the positions of individuals throughout the simulations. When the mating distance was small, individuals tended to group together and move cohesively throughout the landscape (as shown in Fig. 4). As the mating distance increased, the population broke into discrete clusters which appeared to “repel” each other. Varying the competition distance had little effect on spatial clustering.

We next examined how the dispersal, competition and mating distance affected a set of summary statistics for the genealogies (Fig. S1). We computed Sackin’s index, as well as two measures of diversity: the average number of pairwise differences (Tajima’s estimator of diversity) and the number of segregating sites for each of the trees, as described in Methods section 5.1.4.

The average number of pairwise differences decreased with the dispersal distance, and the number of segregating sites showed the same pattern. This suggests that limited dispersal range preserved diversity in the population, although it appears to be inconsistent with the well-known Wahlund effect, the decrease in diversity brought about by population structure (the Wahlund effect, Wahlund 1928). Interestingly, increasing the mating distance instead led to an increase in diversity and the number of segregating sites, which is instead in agreement with the Wahlund effect. The average number of pairwis and number of segregating sites showed no clear pattern with increasing competition distance.

Furthermore, the Sackin index exhibited a reduction with increasing dispersal distance, while it remained constant when altering mating and competition distances. Sackin’s index, a measure of tree balance, is defined as the sum of the number of ancestors for each tip of a tree (Lemant et al. 2022). A higher Sackin index signifies a less balanced tree, indicating that certain clades tended to give rise to more descendants than others. Consequently, this pattern suggests that short-range dispersal introduced some imbalance into the branching structure of the genealogies.

### 3.2 Modelling dispersal patterns

Inspired by these observations, we developed a theoretical model of parent-offspring distances combining *σ* and the scale of mating, given a mode of dispersal where distances were drawn from a Gaussian distribution (which here we term “Brownian”, as described in Methods section 5.1.1) using the uniform model of mate choice implemented in *slendr*. This also represents a more general example of a species for which mate choice and dispersal distances are not drawn from the same distribution, or at the same scale.

The distribution of parent-offspring distances is an equally weighted mixture of dispersals from a “gestating” parent *p*1 and a non-gestating parent, *p*2. If the parent-offspring distance is *y*, its density given a dispersal distance parameter *σ* and a mate choice radius *r*_*b*_ is

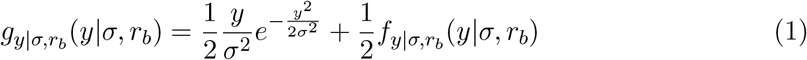

The first term reflects the density given a standard Rayleigh distribution (between the gestating parent and its offspring) with scale *σ*, while the second term models the distance between the non-gestating parent and the offspring.

If we assume a uniformly distributed mate choice radius, then the density function of the distance between the non-gestating parent and the offspring is given by

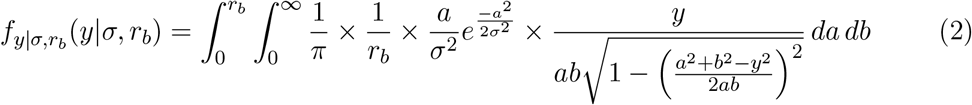

Where *a* is the distance between the gestating parent (*p*1) and its offspring, and *b* is the distance between parents. This derivation is elaborated in the Methods section 5.3. We verified that these equations matched the simulated distances (Fig. 5) across the parameter range we examined.

**Figure 5:**
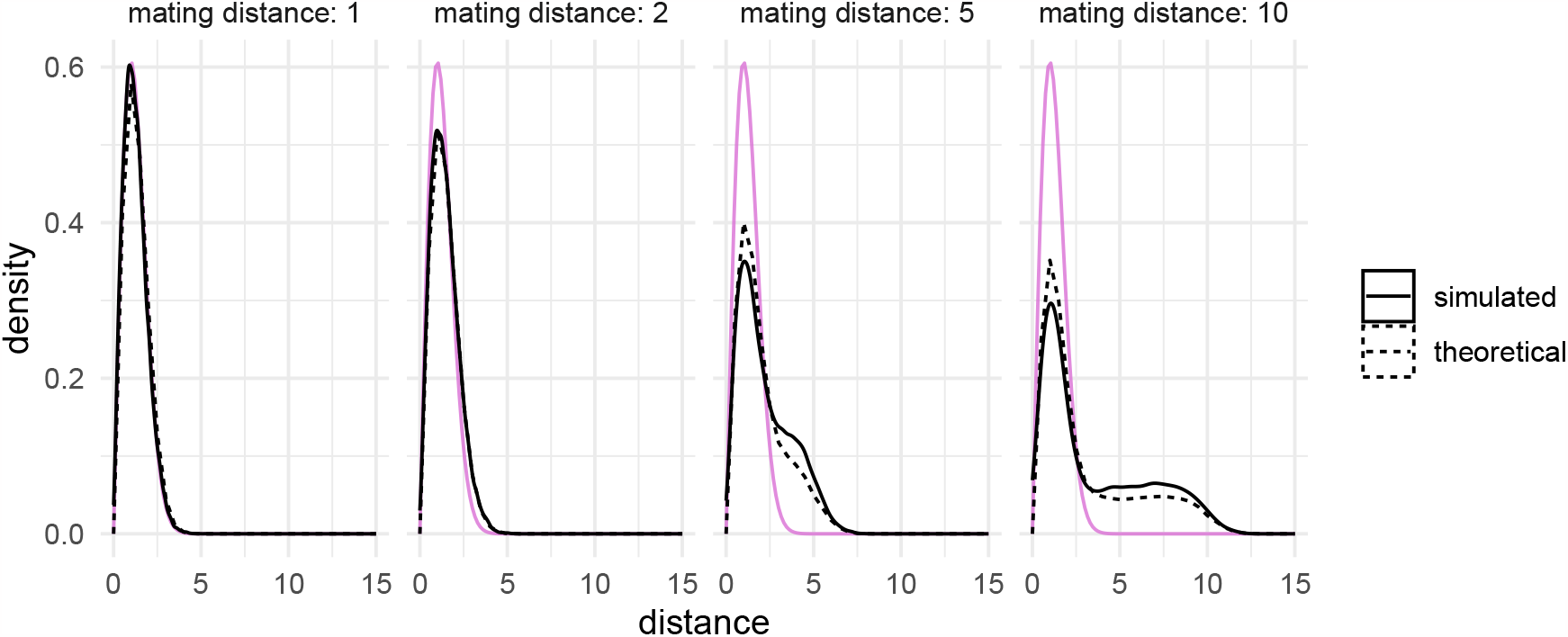
Modelling dispersal and mate choice. The distribution of parent-offspring distances against the theoretical distribution given in Equation (1) (dashed lines) and the Rayleigh distribution (purple). There was a close match between the theoretical and simulated distributions across a range of mating distances. As the mating distance increased, the distributions acquired a flat shoulder compared to the Rayleigh distribution, arising from long father-offspring dispersals. In all simulations, the dispersal distance parameter *σ* was 1 unit and the competition distance was 1 unit.

If the mate choice distance is instead modeled more simply as a Rayleigh distribution (see Methods section 5.3), the density function between the offspring and the (unknown) parent can be analytically solved:

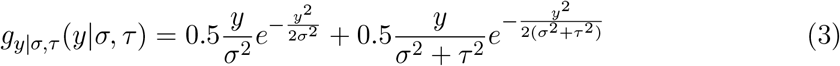

where *τ* is scale of the Rayleigh distribution governing the mate choice distance.

This formulation also leads to a simple result for the mean parent-offspring distance. Since the expected p1-offspring distance is 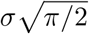 and the expected *p*2-offspring distance is 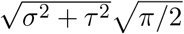, the expected parent-offspring distance is 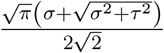. If we were to measure the distances along branches of a genealogy, we would eventually expect to see generation-scaled distances follow a Gaussian with mean 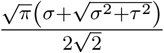. This may be interpreted as a many-generation “effective” dispersal distance parameter.

### 3.3 Estimating dispersal distances from spatially tagged genealogies

Finally, we sought to test how accurately *σ* could be estimated, given a perfectly inferred spatial tree sequence. Under a Gaussian mode of dispersal (what we term “Brownian”), and negligible mate choice distance, the maximum likelihood estimator of *σ* is

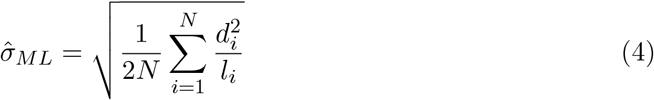

where the index *i* denotes each of *N* branches in a genealogy, with geographic distance *d*_*i*_ and branch-length in generations *l*_*i*_ (see Methods section 5.4). It is worth noting that our method of estimation is naïve, since it ignores the fact that branches are shared between pairs containing the same individual — indeed, we actually maximize the composite likelihood, rather than the full likelihood (as instead is used in Osmond and Coop 2021, where covariance between pairs is appropriately taken into account). However, with enough data, the maximum likelihood estimate of *σ* should be the same in both cases, and we use this as a simple bench-mark approach.

We sampled 100 genomes across 5 simulation replicates from a population of size *N* = 2, 000, and set the mating distance to be small (0.2 units) to minimize its effect on dispersal. We first obtained maximum likelihood estimates of *σ* from the set of all parent-offspring distances. We next emulated a situation where the geographic positions of tree tips and internal nodes are known, but those of the individuals along lineages in the tree are not known (labelled “*simplified*” in our plots). Lastly, to mimic a more realistic scenario, we extracted the distances between all pairs of tips, which corresponds to a situation where only present-day individuals have a known location (“*tips only* “). The results are shown in Figure 6.

**Figure 6:**
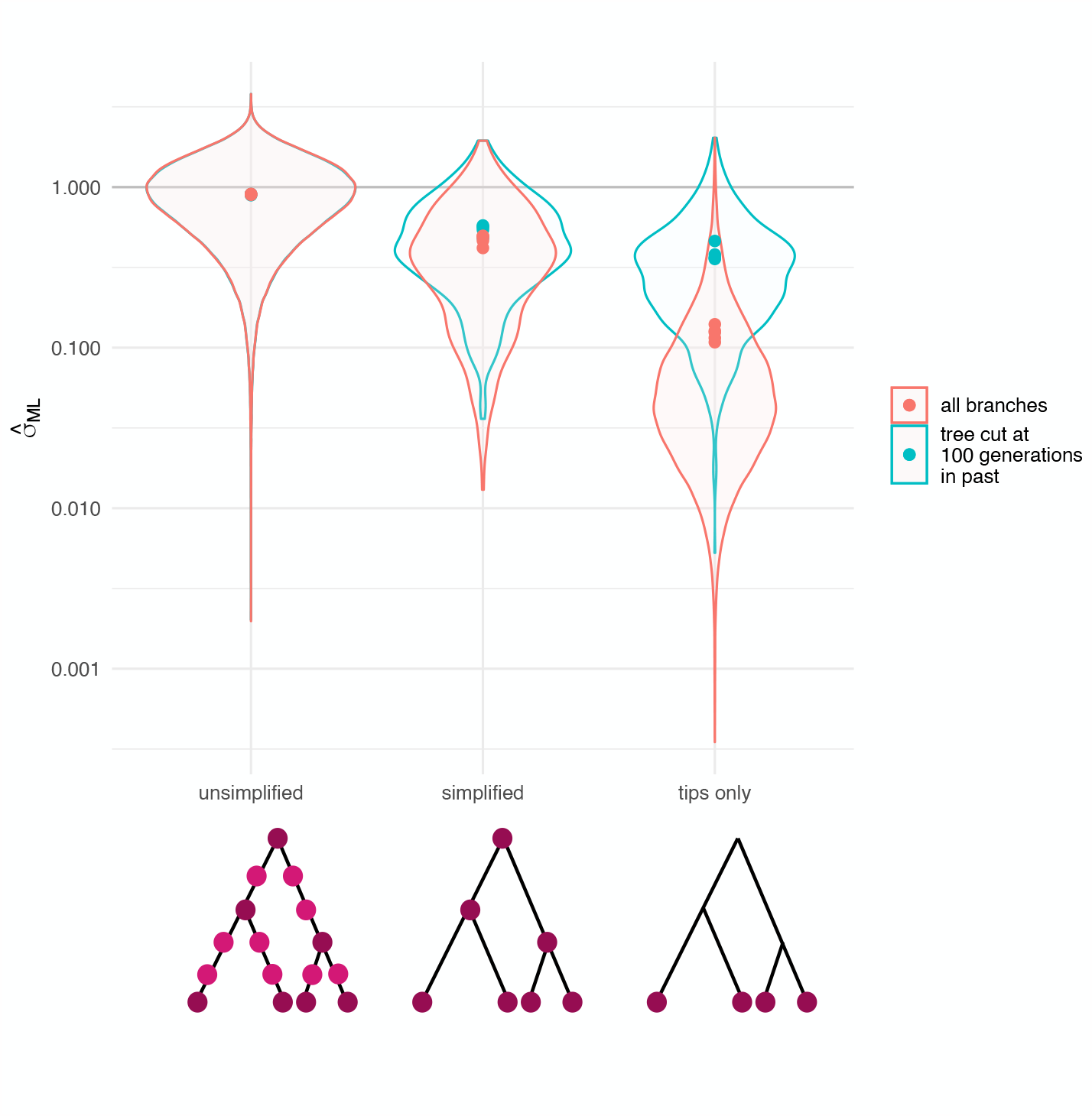
Estimating the dispersal distance under a Brownian dispersal kernel. Each dot is the ML 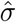, from each of 5 simulation replicates. The violin plots show all branch-wise 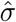 values and the grey lines show the true *σ*, 1 unit. The diagrams below illustrate the lineages used in each case. Excluding older branches, as in Osmond and Coop 2021, increased the estimated dispersal distance for the simplified tree and the tip-only distances. We suggest that this is because more ancient, longer branches in the genealogy are biased due to limited habitat size. In each case, the mating distance was 0.2 units and the competition distance was 0.2 units.

While the estimates of *σ* from the full set of parent-offspring distances were accurate, the estimates from longer tree branches generally were smaller than the true parameter. To investigate whether limited world size was responsible for this observation, we adopted the approach detailed in Osmond and Coop 2021 and eliminated branches which were more than 100 generations old. In the “*simplified*” and “*tips only*” case, this amounted to retaining sub-trees for which the tMRCA lived less than 100 generations in the past. Pruning the trees caused the distribution of branch-wise distances to more closely resemble that of the simplified trees, and correspondingly caused an increase in the estimated *σ*. This suggests that distances accumulated over long branches in a given genealogy tended to be shorter than expected: a phenomenon probably caused by the fact that long-range dispersal is limited in a finite habitat. This pattern was consistent across a range of dispersal distances (Fig. S2).

We also tested whether assuming an incorrect dispersal kernel could affect estimates of *σ*. This might be applicable in a situation where, for example, a population follows power-law dispersal, but we assume parent-offspring distances to be Gaussian and attempt to estimate the variance parameter. Another way to interpret this, is to estimate the net effective dispersal parameter which results from a Cauchy *DF*. To mimic this situation, we simulated under a mode of dispersal where a random angle was drawn from a uniform distribution and a distance from a Cauchy distribution with scale and location of 1 unit. The Cauchy distribution is more heavy-tailed compared to a Rayleigh distribution with the same scale. In agreement with this, the estimated *σ* was larger than the true parameter (Fig. 7). We also varied the scale of mate choice to see what synergy large mating distances might have with a misspecified dispersal kernel. As expected, the estimates of *σ* increased as with the scale of mate choice. Interestingly, there appeared to be a steeper increase in 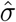 with mating distance when the *DF* was Cauchy.

**Figure 7:**
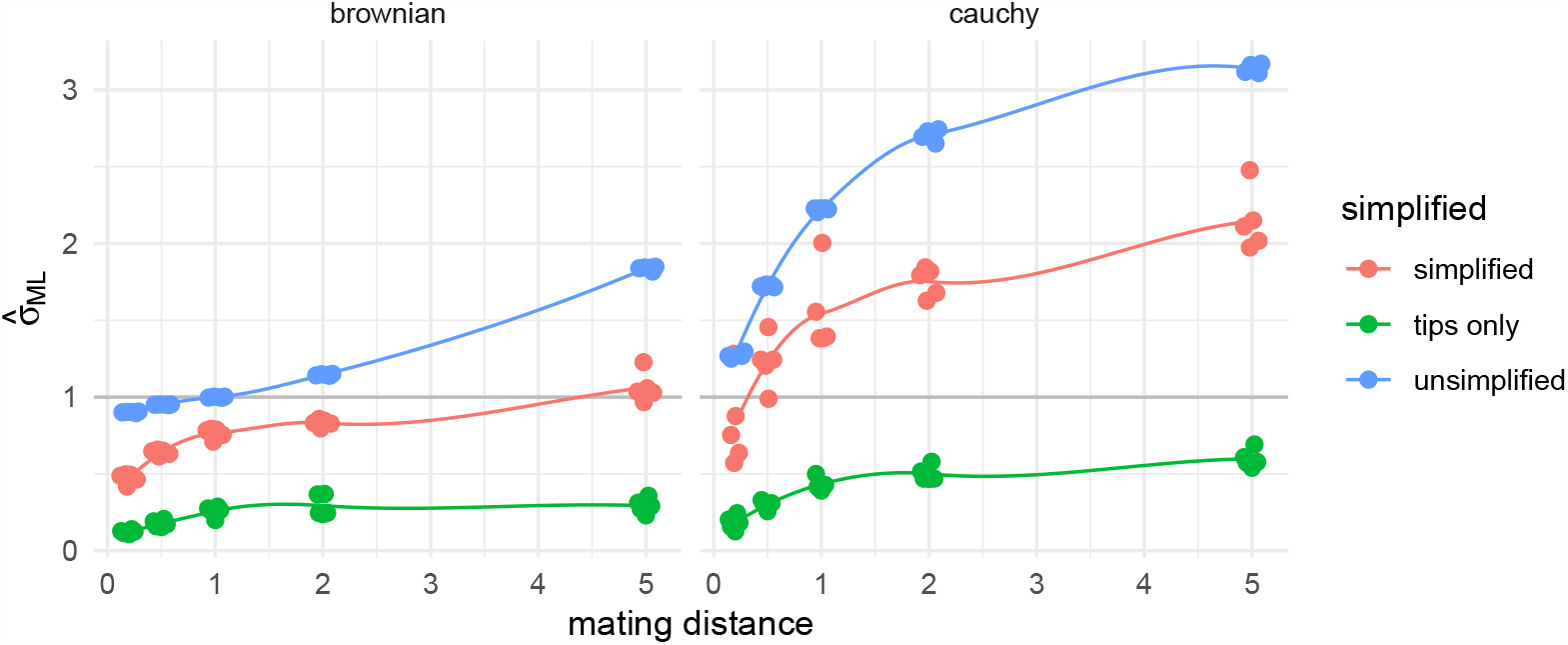
Estimating the dispersal distance under a misspecified model. Left: increasing the mating distance increases the effective dispersal distance. Right: in these simulations, the dispersal function was Cauchy with scale and location 1 unit, but we naively used maximum likelihood estimator of *σ* for the Brownian mode of dispersal. In this setting, increasing the mating distance led to further inflation of 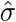. The dots show the result of each of five replicates, and the lines are smoothed rolling means.

## 4 Discussion

In this study, we explore the effects of three important ecological parameters (dispersal distance and distribution, competition distance and mate choice distance) on the geographic distances captured in a geographically tagged genealogy.

We show that altering the kernel of parent-offspring dispersal can have strong effects on the diffusion captured within a genealogy, and in particular on the tails of the realised parent-offspring distance distribution. For example, the Cauchy distribution, which is a text-book example of a “heavy-tailed” distribution, did indeed produce a greater proportion of long-distance dispersals.

There was some difficulty in choosing a common parametrization for these dispersal distributions, especially since *slendr* implements two different mechanics of parent-offspring dispersal (one where a random distance and angle are chosen, and another where latitudinal and longitudinal distances are chosen, see Methods section 5.1.1). We suggest that a pragmatic solution for the sake of simulation might be to encode a dispersal distribution where the height of the tail may be controlled more directly. An example may be the Pareto distribution, where the tail probability is particularly sensitive to the shape parameter, and does not directly depend on the variance (in contrast to, for example, the normal distribution).

The mate choice radius caused distinctive patterns in the distribution of a population within its landscape. In particular, close-range mating led to clustered groups of individuals, which may be a practical nuisance to simulation users, and lead to unwanted geographic structure. We suggest that this is the same phenomenon described in Felsenstein 1975. As Felsenstein describes, the intuition behind this behaviour is that, when either mate choice or dispersal distances are small, individuals each seed a “clump” of descendants. Due to the constraint of constant population size, several of these clumps are destined to die out. The small mating distance forbids mating between these clumps, so the remaining ones become larger and further apart. This is particularly cumbersome because relatively small mating distances are required for the average parent-offspring dispersal to match p1-offspring dispersal. Although not possible in the most recent version of *slendr* (Petr et al. 2022), allowing for less generally constrained simulations with fluctuating population size might alleviate these factors. However, this would require the development of dedicated software for the analysis of tree sequences produced by such dynamics (known as “non-Wright-Fisher”) in *slendr*.

We also observed that the distances within a genealogy increased dramatically if the scale of mate choice was large. Mating is often not explicitly modelled — yet the step of mate allocation is essential in forwards-in-time, agent-based genetic simulators such as *slendr* and *SLiM*. Furthermore, the dynamics of mate choice and parent-offspring dispersal may differ starkly in natural populations: for example, the same model of dispersal may not apply to the dynamics with which pollen and seeds spread. Our results support that this is an important parameter, and absorbing mating and parent-offspring dispersal dynamics into one step may not always be appropriate.

Aside from changing the distances in the genealogy, the scale of mate choice also changed the shape of the distribution of parent-offspring distances. To illustrate a case where this may be modelled, we described the theoretical distribution of parent-offspring distances under uniform mate choice, and found a close match between the this and simulated distances. The natural next step would be to use these results in an inference framework, by deriving analytical solutions for the maximum-likelihood or method-of-moments estimators for the dispersal and mating distances.

Rather than the theoretical dispersal distance itself, a parameter that may be more liable to inference is an effective dispersal distance parameter, which incorporates both the mate choice and dispersal processes. The distance between parents and offspring over many generations should follow a normal distribution in the limit of infinite generations, due to the central limit theorem. Therefore, if we were to take the distances along branches of a phylogeny and scale them by the respective number of generations (as inferred from genetic data), the distribution of distances would approach a Gaussian distribution, centred around this effective dispersal distance. Specifically, this is an equally weighted mean of the expected distances of the offspring from either parent (see Methods section 5.3). For example, in the Methods section 5.3.1, we show that under a model with Gaussian dispersal (with scale *σ*) and mate choice (with scale *τ*), this effective dispersal distance can be easily calculated as 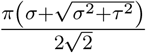.

This compound parameter is in effect what is estimated when mate choice dynamics are not explicitly modelled in phylogeographic studies. We therefore motivate distinguishing between spatial models intended for few generations, where the stages of mating and dispersal should be treated as distinct, from those for phylogenetic time-scales, where they may be absorbed into one parameter. We also note that, over long time-scales, dispersal was limited by finite population ranges. In our results, this led to estimates of the mean dispersal distance which were smaller than expected, illustrating that deep coalescent branches should only be used with caution for inference, as illustrated by Osmond and Coop 2021.

This study has focused on single-locus genealogies, which is comparable to studying approximately independent genealogies from a tree sequence. Such an approach, followed for example in Osmond and Coop 2021, greatly reduces the computational burden of analysing the full tree sequence, yet retains the ability to uncover variation in dispersal and geographic ancestry across the genome. However, we expect that ignoring the correlation structure which exists between trees in a tree sequence leads to some loss of information — specifically, in a fully annotated tree sequence, it is possible to identify nodes which are shared between trees. This information could be used, for example, to constrain the positions of shared internal nodes based on information coming from several trees. We note that, since *SLiM* and *slendr* are able to run spatial simulations of recombining genomes, they might be valuable tools to begin to investigate how much information is lost when we “thin” tree sequences.

Another aspect of complexity which we have not investigated is the bias which might arise from using estimated genealogies, rather than known ones. There is recent evidence that currently available methods (*Argweaver, Relate* and *tsinfer + tsdate*) tend to under-estimate the time of deep coalescences, and vice versa (YC Brandt et al. 2022). This is a form of a well-known phenomenon in phylogenetics called “long branch attraction”. We expect that would lead to biases in inferences of dispersal (longer-range than reality towards internal nodes, and shorter than expected at the tips). Again, this could be aptly studied in *slendr* by *post-hoc* adding mutations onto the simulated genealogies, and adding a genealogy estimation step to the analyses.

In cases where we are interested in untangling the mating and dispersal distances, uniparentally inherited genetic material could be of use. Mitochondrial DNA only moves via mother-offspring dispersals, the direct manifestation of the dispersal function (when the mother is *p*1). Conversely, the Y-chromosome always moves according to a convolution of mating and dispersal distances. Comparing their respective rates of diffusion could help us identify cases in which the between-parent distance might be masking the underlying mother-offspring dispersal dynamic.

At the moment, *slendr* is not able to model sex differences. Yet, mother-offspring dispersal and mate choice may span different scales if dispersal is strongly sex-biased. Theoretical results across a range of animals suggest that this is the case when the limiting resource differs between males and females (Li and Kokko 2019). In line with this, field observations and genetic data have pointed to a breadth of matrilocal and patrilocal behaviours across animal species (for example Liebgold, Brodie III, and Cabe 2011; Oota et al. 2001; Schubert et al. 2011). These sex-biased processes might be an intriguing direction for further investigation.

Another exciting direction for further study is selection. A positively selected allele will often have more descendants than a neutral one, resulting in excess branching. This means that positively selected loci, and genomic regions in linkage disequilibrium with them, are expected to have more descendant lineages which can explore space and travel faster than neutral ones. This result is similar to Fisher’s travelling wave model, where the velocity of spread is proportional to the square root of the selection coefficient (Fisher 1937; Muktupavela et al. 2021; Steiner and Novembre 2022). For the purpose of inference, we often assume that the coalescent branching process and geographic location are independent (although this is not the case, see Wilkins and Wakeley 2002). How far do we deviate from this assumption, for example, when selection pressures are local?

Overall, it is clear that accurately modelling the dispersal of a given species may require sound understanding of a variety of ecological parameters. From our simulations, we observed that geographic distances captured within a geographically tagged genealogy captured these compound effects. These are not yet theoretically well-understood, and may become confounding factors in joint analyses of geographic space and genetic diversity. Simulations will be key to approaching these issues.

## 5 Methods

### 5.1 Spatial simulations

We used the software *SLiM* (Haller and Messer 2019) via its R interface *slendr* (Petr et al. 2022) to simulate populations in space and time.

Generations were discrete and non-overlapping, and there was no modelled age structure or sex-based differentiation. We chose to keep populations at a constant size in order to focus on fundamental aspects of dispersal without confounding effects from demographic size changes.

At each generation and for every individual, the program counted the number of neighbours within a radius of the competition distance (let this be *n*). Then, the fitness was down-scaled by this number to model competition for resources (fitness ∝ 1*/n*).

Individuals were chosen randomly, weighted by their fitness, to be the parents of the next generation. Pairs of mates were chosen within a radius of the mating distance, with uniform probability. Within each of these pairs, one parent at random was set to be *p*1, which is sometimes called the “gestating parent”. However, note that this is purely a label — it may also be that *p*2, whether it be the mother or the father, migrates to *p*1’s position to raise the offspring.

In this set-up, the location at which individuals mate is also that at which their fitness is evaluated. These are the coordinates recorded in our simulations. This means that *p*1 − *o* displacement can be seen as the net of parents moving to have the offspring, and the migration over the offspring’s lifetime from their birthplace to their mating location.

In *slendr*, a user specifies a model and its parameters. These are passed to a *SLiM* backend, which executes the simulation. After this, among the data which can be recovered from a simulation are the locations of all individuals, the times at which they lived and the phylogeny and pedigree connecting them.

#### 5.1.1 Encoding dispersal

We simulated under several modes of *p*1-offspring dispersal, coming under two categories:

1. Angle-distance dispersal: in these, the absolute distance is controlled by a given distribution. An angle is drawn randomly from a uniform distribution between 0 and 2*π*, and a distance *d* was drawn from one of the following distributions:
  - *Uniform*: the p1-offspring distance is uniformly distributed between 0 and *σ, d* ∼ *U* (0, *σ*). The mean absolute distance is *σ/*2 and the variance is (1*/*12)*σ*^2^.
  - *Half-Normal* : the p1-offspring distance is Gaussian distributed, with mean 0 and variance *σ*^2^. When a distance is below zero, the offspring is effectively ejected backwards. The mean of the resulting folded normal distribution (specifically, a half-normal) is 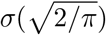 and the variance is *σ*^2^.
  - *Exponential* : the p1-offspring distance is exponentially distributed, with rate parameter 1*/σ, d* ∼ *Exp*(*σ*). The mean is *σ* and the variance is *σ*^2^.
  - *Cauchy* : the p1-offspring distance is Cauchy distributed, with location 0 and rate parameter *σ, d* ∼ *Cauchy*(0, *σ*). The mean and variance of this distribution are undefined.
2. *Brownian*: here, the axial distances are controlled. Random distances in the *x* and *y* dimensions (*d*_*x*_ and *d*_*y*_) are each drawn from a Gaussian with mean 0 and variance *σ*^2^, *d*_*x*_ ∼ *𝒩*(0, *σ*^2^), *d*_*y*_ ∼ *𝒩*(0, *σ*^2^). This means that the absolute distance then follows a Rayleigh distribution with scale *σ*, which has mean 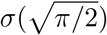 and variance 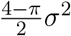. This mode is named “*Brownian*” due to its conceptual relation to a 2-dimensional Brownian motion.

#### 5.1.2 Tree recording and manipulation

We simulated a single locus in order to focus on fundamental geographic dynamics which act on single trees. After a simulation run, we retrieved the simplified and unsimplified trees. Simplified trees, which are the same as standard coalescent trees, consist of nodes representing coalescence events, and edges connecting them. These edges implicitly record many individuals. In contrast, an unsimplified tree records all individuals along edges. Such a tree is useful to directly observe the dispersals which occurred at every generation along a long branch. We processed and analysed these via the *slendr* interface to the *tskit* library (Kelleher, Thornton, et al. 2018). *tskit* is a powerful framework for storing and manipulating trees and tree-sequences with close-to-optimal space usage. We also converted these trees to the “phylo” R object class, which allowed us to analyse them via the phylogenetics package *ape* (Paradis and Schliep 2019).

#### 5.1.3 Geo-spatial analyses

*slendr* integrates with the spatial package *sf* (Pebesma et al. 2018), and this allowed us to extract a variety of spatial features from the trees, including the positions of individuals, the vectors connecting nodes and the distances between them.

#### 5.1.4 Computing tree statistics

We computed the normalized Sackin’s index using the R package, *apTreeshape* (Bortolussi et al. 2006). In order to compute the number of segregating sites, we used *slendr* ‘s *ts_segregating* function in “branch” mode. To compute the diversity (the average pair-wise difference between sequences), we added mutations to the genealogies post-hoc with *ts_mutate*, and then applied the *ts_diversity* function.

### 5.2 Statistics and Plotting

We calculated statistics in base *R*, as well as with the packages *VGAM* (T. W. Yee, M. T. Yee, and VGAMdata 2022) and *moments* (Komsta and Novomestky 2015). We evaluated numerical integrals in Mathematica (Wolfram 1991). We produced plots with *ggplot2* (Gómez-Rubio 2017) and auxiliary packages.

### 5.3 Derivation of the probability density of the distribution of parent-offspring distances

A diploid individual carries two genome copies, each inherited from a parent. These have a distinct genealogy and in any given tree, we follow the movement of one of these copies through individuals over time and space. We can therefore break down the dispersals which occur in one generation into two categories:

1. Genetic parent is the “mother”, *p*1. We observe *p*1-offspring dispersal, (which in *slendr* is directly encoded).
2. Genetic parent is the “father”, *p*2. We observe a convolution of *p*1-offspring dispersal and the *p*1 − *p*2 distance.

We can draw a triangle which connects both parents and offspring, as shown in Fig. 8. In case (1), we observe side *ã*. In case (2), we observe side 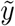. 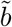 is the distance which separates the two parents, and the angle between sides *ã* and 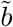 is 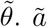 ∼ *Rayleigh*(*σ*), if we have *Brownian* dispersal. Since in *slendr*, parents are chosen with uniform probability from a specified radius *r*_*b*_ (the mating distance), 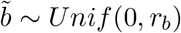 where *r*_*b*_ is the mating distance. The angle between these sides is free to range between zero and *π*, so 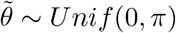.

**Figure 8:**
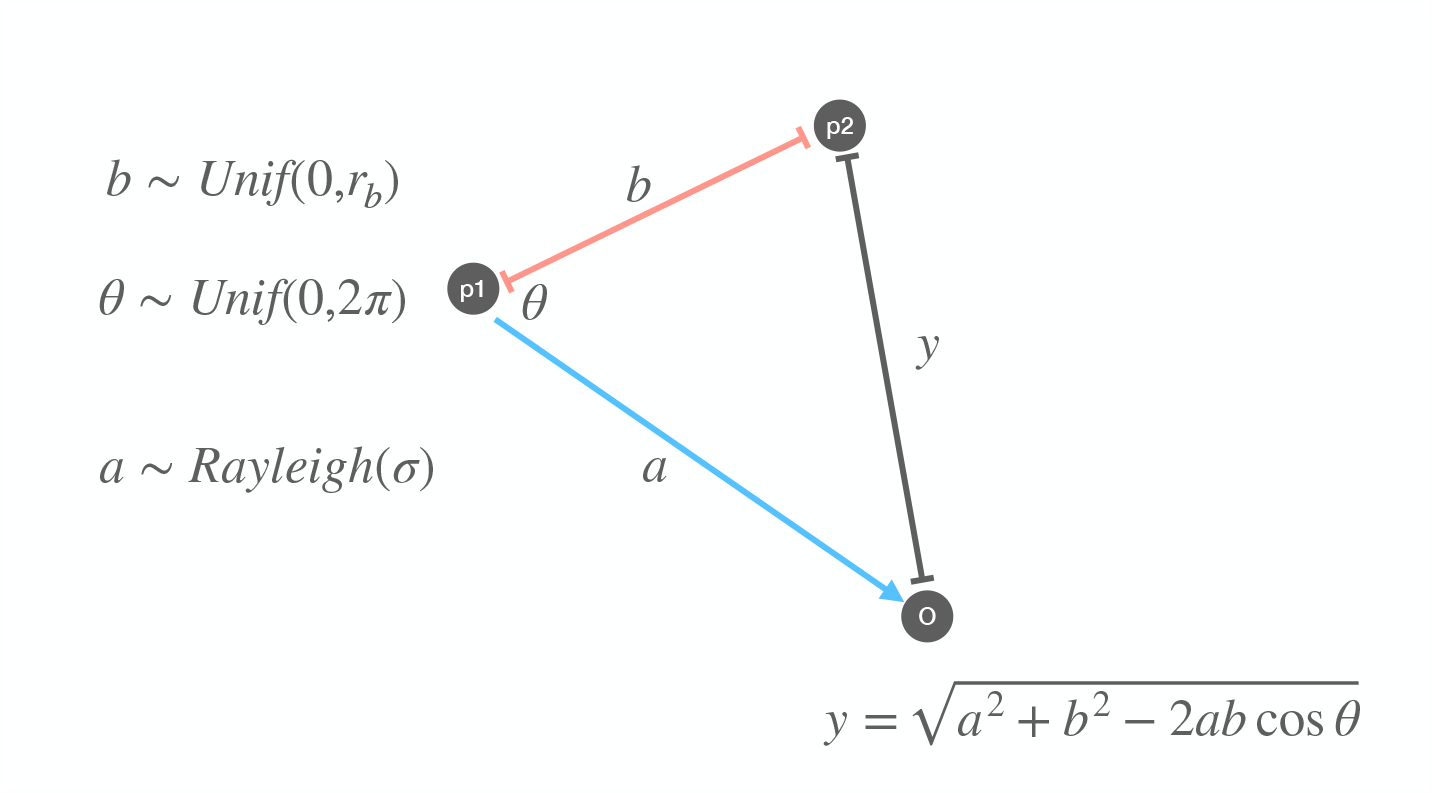
A schematic of parent-offspring dispersal. When we observe dispersal from *p*2, the observed parent-offspring distance (*y*) is a convolution of the distance between *p*1 and *p*2 (*b*, in red), and the dispersal between *p*1 and the offspring (*a*, in blue). The cosine rule gives us an expression for *y* in terms of *a, b* and the angle between them *θ*. If we know the probability distributions of *a, b* and *θ*, we can obtain that of *y* via a change of variables.

We can calculate the length of the side *y* from *a, b* and *θ*:

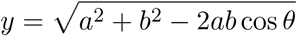

We aim to derive the probability density function (pdf) of *y*, using the pdfs of *a, b* and *θ*. This can be achieved with a change of variables:

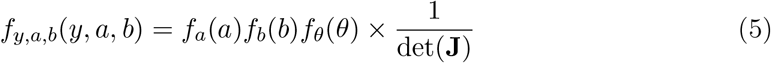

**J** is the jacobian matrix of partial derivatives:

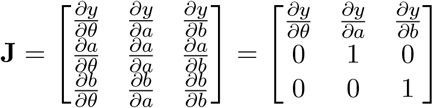

The determinant of this matrix is

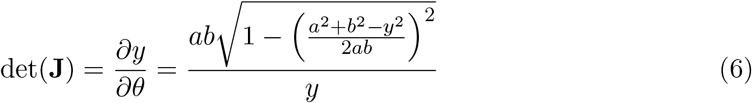

Which goes back into equation (5):

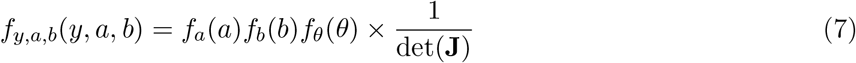

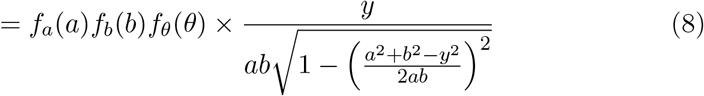

This is the joint pdf of the three sides of the triangle. Now, we integrate out the parameters *a* and *b* in order to get a fully marginalised *f*_*y*_.

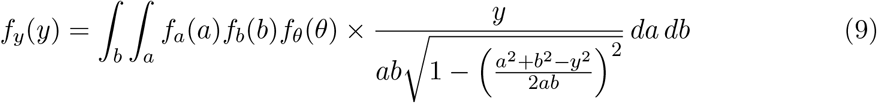

This holds for any distribution of *a* and *b*. Let’s consider the case where *a* is Rayleigh distributed (as it is under the *Brownian* mode of dispersal), and mate choice is random within a radius *r*_*b*_ (as encoded in *slendr*). *θ* and *b* are uniform random variables, so have a constant probability of 1*/π* and 1*/r*_*b*_ respectively. We also know that *a* has a Rayleigh pdf of 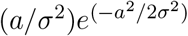. Replacing these in the function above:

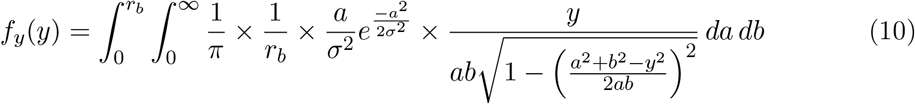

This is the fully marginalised pdf of *y*. This integral is challenging to solve analytically, but we can obtain the approximate shape of the pdf by numerical integration.

Finally, we can write out the pdf of the distance between a randomly chosen parent and its offspring. Let’s call this pdf *g*_*y*_(*y*). With probability *P* = 0.5, the parent is the mother (*p*1) and *y* simply follows a Rayleigh distribution with scale *σ*. When the genome is inherited from the father (*p*2), which again occurs with *P* = 0.5, the pdf of *y* is the distribution shown above. This leads to the final pdf *g*_*y*_(*y*) of the parent-offspring distance,

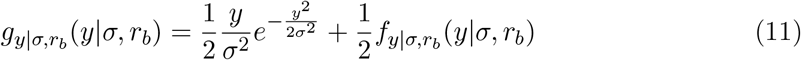

From this expression, we can obtain any moment of the distribution. The expectation of the distance *y* is:

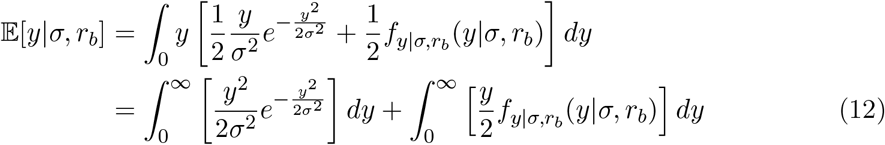

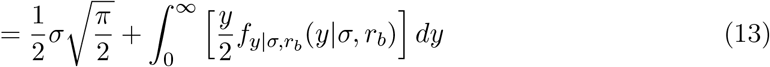

which is a half-weighted average of the distance expected from the parent-offspring distance kernel, and from mate choice.

#### 5.3.1 A simpler model with Gaussian mate choice

There are simple scenarios that lead to a more analytically tractable pdf. For example, let us suppose that the distance between parents is also generated in a similar way to *Brownian* dispersal, from independent normal distributions in *x* and *y* dimensions with variance *τ* ^2^. In this case, the father-offspring distance in each dimension is a sum of two Gaussian random variables and is itself normally distributed with variance *σ*^2^ + *τ* ^2^. This gives rise to a Rayleigh distribution with scale 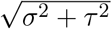 for the norm of the distance, *y*. In that case, the final pdf is then:

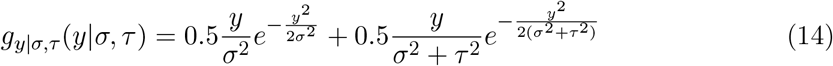

As noted in Battey, Ralph, and Kern 2020, if the scale of dispersal and mate choice are the same (if *σ* = *τ*), the spatial diffusion process becomes Gaussian with an overall variance 3*σ*^2^*/*2.

### 5.2 Maximum likelihood estimation of *σ*

When the mating distance is small, and dispersal is “Brownian”, distances in latitude and longitude at each generation are drawn from independent 𝒩(0, *σ*^2^), and the dispersal over many generations may be modelled as a Brownian motion. Given a genealogy with *N* branches *i*, of length *l*_*i*_ and geographic distance *d*_*i*_, the log likelihood of the distances is

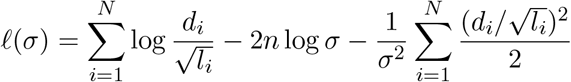

Here, we have divided each branch distance *d*_*i*_ by 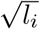 to account for multi-generation branches. The absolute distance should increase proportionally to the square root of the number of generations, since dispersal is Gaussian in two dimensions.

The gradient of the likelihood function with respect to *σ* is

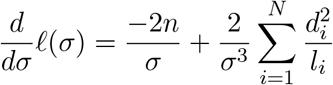

The maximum likelihood estimator of *σ*, which solves 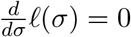, is given by

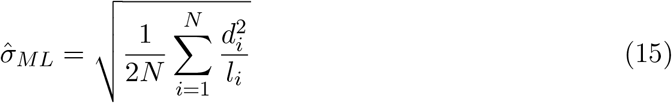

We may also wish to survey how each branch is contributing to the estimate. Since 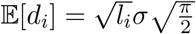, we define 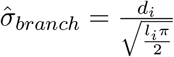.

## 5.5 Code availability

The functions used (which are not included in *slendr* or other packages) are available as an R package *treesinspace* (https://github.com/mkiravn/treesinspace/). We include all relevant scripts, with which the simulations and plots included may be reproduced.

## 5.6 Acknowledgements

We thank members of the Racimo group and the Novembre group for valuable comments on the manuscript, as well as Thomas Bataillon (Aarhus University) and John Novembre (University of Chicago).

FR is supported by a Villum Young Investigator Grant (project no. 00025300), a Novo Nordisk Fonden Data Science Ascending Investigator Award (NNF22OC0076816) and by the European Research Council (ERC) under the European Union’s Horizon Europe programme (grant agreements No. 101077592 and 951385). MP is supported by a Lundbeck Foundation grant (R302-2018-2155) and a Novo Nordisk Foundation grant (NNF18SA0035006) given to the GeoGenetics Centre.

## 6 Supplementary Material

**Figure S1:**
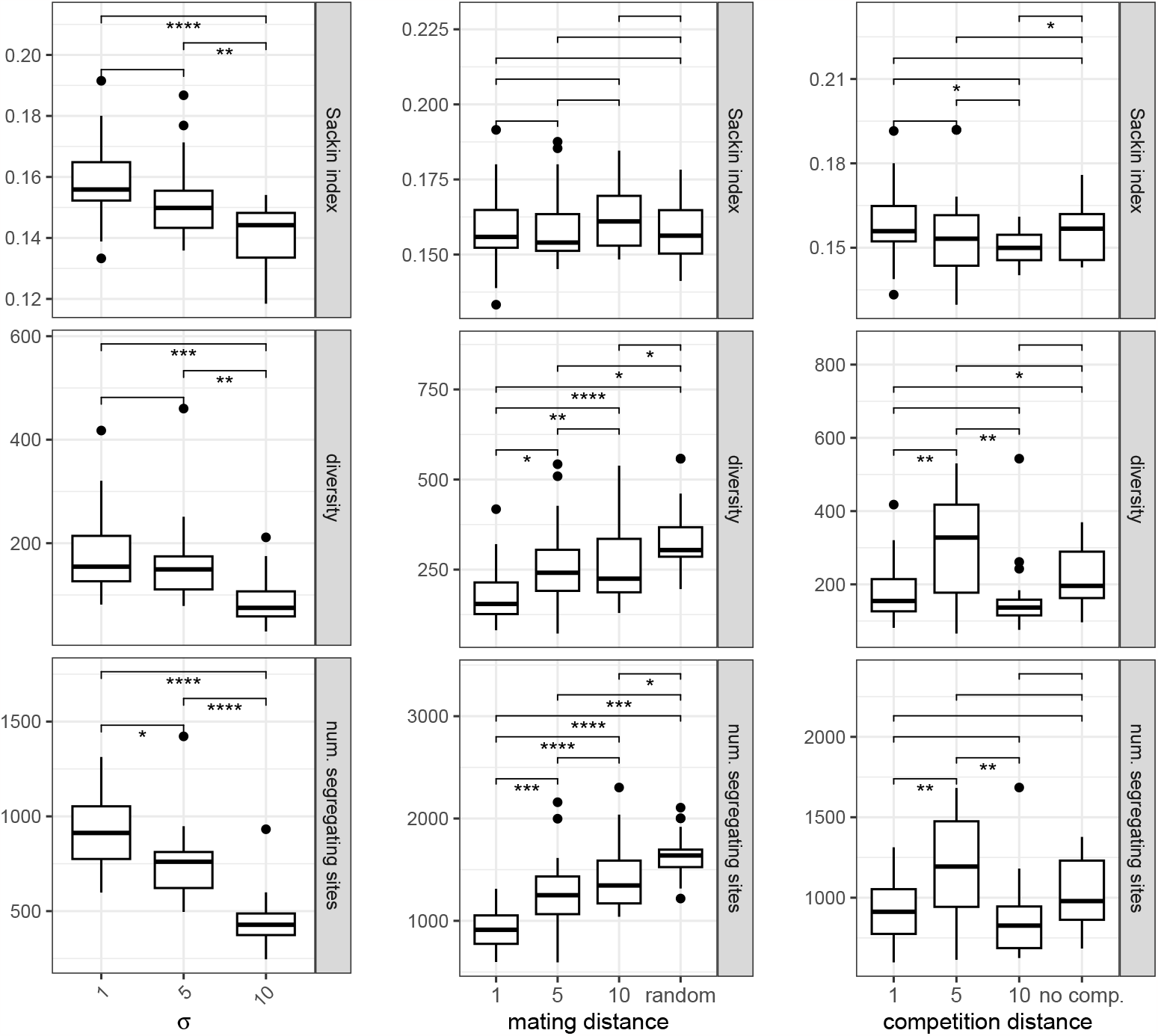
The effect of dispersal, mate choice and competition on tree-based statistics. Each column shows the effect of increasing one parameter, with the others kept constant at 1. Stars show the level of significance of a two-sided t-test. The diversity was calculated as average pair-wise difference between sequences; the Sackin index is the sum of leaf depths for a given tree and reflects tree balance (less balances trees have a higher Sackin index). We ran 20 replicates of a simulation with 100 individuals with Brownian dispersal, over 500 generations.

**Figure S2:**
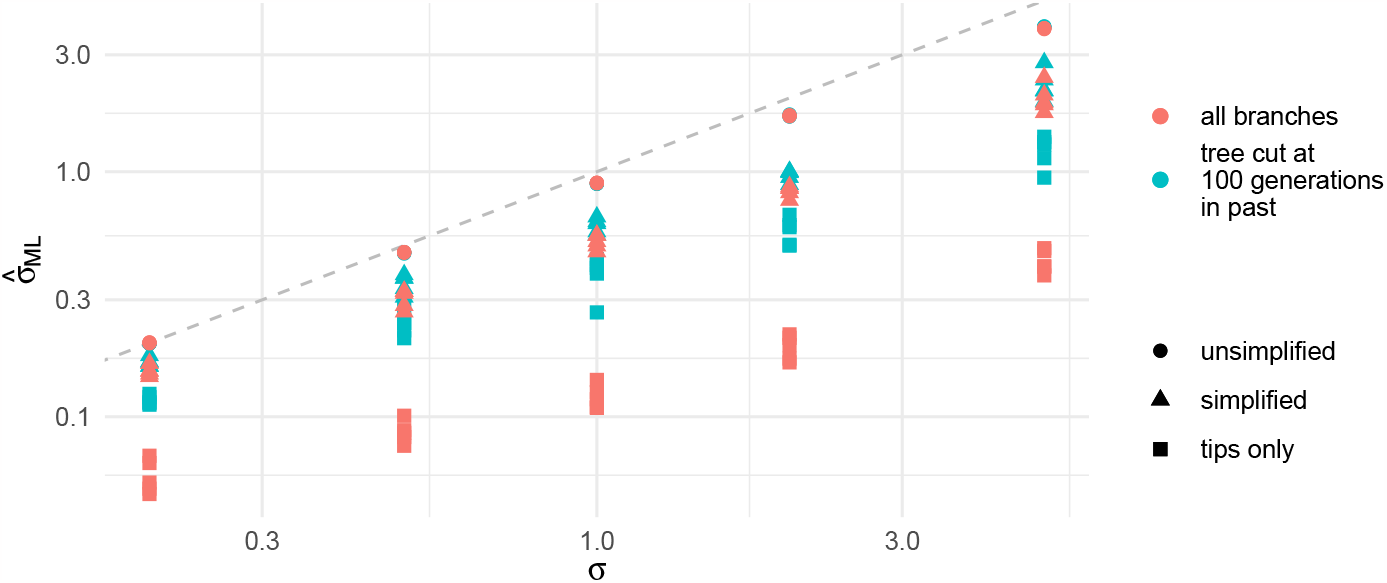
Estimating the dispersal distance with Brownian dispersal, across a range of *σ* values. The grey line shows the true *σ*. We found that the pattern of bias shown in (a) was replicated across the range of *σ* values tested. In these simulations, the mating distance was 0.2 and the competition distance was 0.2.

